# Dysbiotic oral and gut viromes in untreated and treated rheumatoid arthritis patients

**DOI:** 10.1101/2021.03.05.434018

**Authors:** Ruochun Guo, Shenghui Li, Yu Zhang, Yue Zhang, Guangyang Wang, Yufang Ma, Qiulong Yan

## Abstract

**Background:** Rheumatoid arthritis (RA) has been considered to be influenced by bacteria from the oral cavity and gut for many years. Despite potential impact of viruses in RA was mentioned in some studies, specific roles of oral and gut viromes in RA is still unclear.

**Results:** In this study, we observed the viral community variation in the oral and gut samples, performed a comparative analysis of oral and gut viromes in health controls, untreated and treated RA patients, and constructed interaction networks among viruses, bacteria, and RA-associated clinical indexes to address the potential associations between viral community and RA. The results showed that the viromes could be isolated from dental plaque, saliva, and feces samples, among which the saliva having the highest with in-sample diversity. Meanwhile, remarkable variations of viral diversity and composition in the oral (i.e., dental plaque and saliva) virome could be observed in RA patients and healthy controls yet in untreated and treated RA patients, with a relatively low variability in the gut virome. Distraction of viruses-bacteria interaction network was discovered in three sites of RA patients. In addition, some RA-associated oral taxa, including *Lactococcus phage vOTU70, Bacteroides vulgatus, Lactococcus lactis, Escherichia coli, Neisseria elongate*, were correlated to the RA-related clinical indexes.

**Conclusion:** Whole-virome analysis illustrated the potential role of oral and gut viral communities in the development of RA.

## Introduction

To date, the role of the human microbiome in rheumatoid arthritis (RA) was widely studied [1-9]. In the gut, an increased abundance of *Prevotella copri* and decreased abundance of *Bacteroides, Bifidobacterium*, and *Eubacterium rectale* were associated with untreated recent-onset RA [10]. In the oral, the *Porphyromonas gingivalis* was frequently identified as an opportunistic pathogen for RA development[11, 12]. Another whole-metagenome shotgun sequencing-based study revealed an over-expression of *Lactobacillus salivarius* and reduction of *Haemophilus* in the gut as well as dental plaque and saliva of RA patients [2]. Besides, the lung microbiota was also related to rheumatoid arthritis, probably associated with the abundance of some *Prevotella* and *Streptococcus* members in the human respiratory tract [13].

As an important microbial part of the human microbiome, the viral community was also widely considered to be associated with multiple diseases. For instance, HIV infection could lead to an increase of Adenoviridae sequences in human gut [14] and cervical virome[15]. It was also proven that the vaginal viruses are associated with BV-status, serving as an implicit factor in the perturbation of the bacterial community[16]. In addition, Clooney et al. observed the change of gut viral composition with the increased numbers of temperate bacteriophages in Crohn’s disease [17]. Earlier studies found that bacterial communities affect these diseases as well [18-22]. However, as microbes about RA, a disease highly associated with human microbiota, were often identified based on whole-bacteriome analysis, we knew little about the connection of RA and viromes. Notably, the potential impact of virus in RA was widely hypothesized[23-26].One study indicated that respiratory viral infections might bea risk factor for the development of RA[23]. Some studies suggested that *Epstein–Barr virus* (EBV)contributed to the pathogenesis of RA, and an abnormal increase of EBV-infected B cells in the blood, brain, ectopic lymphoid structures of RA patients [24, 26-28]. These findings showed that virus may play an important role in the development of RA. However, the relationships between viral community and RA have not been reported, thus in this study, we investigated the alteration of the viral community and ecological networks of bacteria and viruses in RA patients by whole-virome analysis based on human oral and gut samples.

In this study, we reanalyzed the dataset of Zhang et al.’s study [2], involving a total of 497 samples from three body sites (dental plaque, saliva, and feces) of 102 healthy controls, 104 untreated RA patients, and 64 treated RA patients. We clarified the viral population in human oral and gut communities, and further assessed the potential perturbation of these viromes in untreated and treated RA patients. Meanwhile, we identified interaction networks among viruses, bacteria, and RA-associated clinical indexes in this cohort.

## Results

### Subjects

This study involved a cohort of 266 individuals (164 RA patients and 102 healthy controls) from Shanghai, China. 104 RA patients were new-onset without any medication treatment, while the other 60 patients were treated with multiple drugs. The phenotypic and clinical characteristics of the individuals were introduced in the preliminary study by Zhang *et al*[2].44.7% (119/266) individuals provided both fecal and oral (dental plaque and saliva) samples, while 34.2% (91/266) individuals provided all three samples. Finally, 143 dental plaque (51 healthy controls, 54 untreated patients, and 38 treated patients), 122 saliva (47 healthy controls, 51untreatedpatients, and 24 treated patients), and 232 fecal (97 healthy controls, 94 untreated patients, and 41 treated patients)samples were analyzed. For each body site, the corresponding individuals’ gender, age, and body mass index (BMI) were matched and elaborated in the original study[2].

### Viral population in oral and gut communities

To characterize the oral and gut viral communities, we analyzed a total of 2.72 Tbp high quality non-human metagenomic data (5.48±1.77 Gbp per sample) from 497 samples of dental plaque, saliva, and feces. Metagenomic assembly on each sample generated totaling 2,664,228 contigs (≥5000 bp) (Table S1), of which, 6.1% (n=161,828) were recognized as highly credible viral fragments based on their sequence features and homology to known viral genomes (see Methods).After removing the redundant viral contigs with 95% nucleotide similarity[29], a total of 19,483, 28,123, and 30,805 vOTUs were identified from the dental plaque, saliva, and gut metagenomes, respectively. To investigate the human virome across oral and gut, we combined the vOTUs of three habitats and generated an integrated non-redundance catalog of 48,718vOTUs for further analysis. Based on the assessment by CheckV[30], 15.5% of the vOTUs were high- or medium-quality viral genomes, and the remains were usually low-quality viral fragments or undetermined sequences (Figure S1).55.1% of the vOTUs could be classified into known families. However, only 6% of the vOTUs manifested≥95% nucleotide similarity with available sequenced viruses in the RefSeq database, highlighting a considerable novelty of the oral and gut virome dataset.

Totaling 37,229, 36,803, and 43,170 vOTUs were observed in dental plaque, saliva, and gut metagenomes, respectively. The dental plaque and saliva shared a markedly high proportion (93.6% in dental plaque and 94.6% in saliva) of vOTUs between each other, and they both shared a relatively lower number of vOTUs with the gut virome (85.2% in dental plaque and 84.9% in saliva; Figure 1A). 77.9% vOTUs of the gut virome were also detected in oral samples, and the remaining 22.1% vOTUs were gut-specific. Focusing on the viral diversity in every sample of three habitats, we found that the fecal samples had a significantly high proportion of viral sequences (on average, 14.0%) in their metagenome compared with that of the dental plaque (6.3%) and saliva (9.5%) (Figure 1B left panel). However, the saliva samples had the highest within-sample diversity (estimated by Shannon index) and richness (estimated by the number of observed vOTUs) than dental plaque and fecal samples, while these parameters were lowest in the fecal samples (Figure 1B; Figure S2).

**Figure 1.**
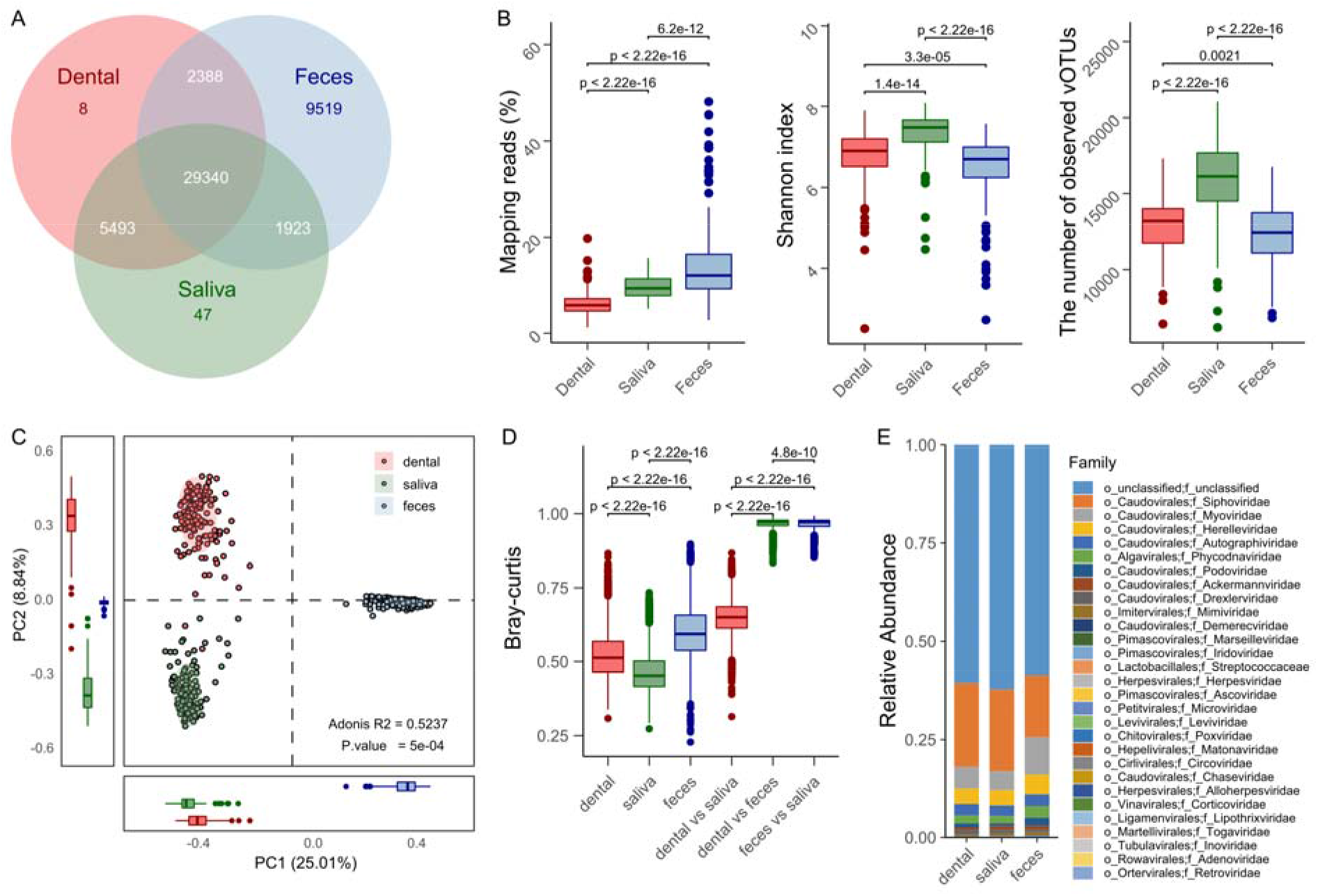
Overview of the oral and gut viromes. **A** Overlap of vOTUs between the dental plaque, saliva, and fecal virome. **B** Comparison of the fraction of reads mapping to all vOTUs and the alpha diversity indexes between the three habitats. **C** Principal coordinate analysis (PCoA) based on the vOTU composition of samples from the three habitats. **D** Bray-curtis distance among samples by body habitat. **E** Relative abundance of viral compositions at the family level of the three habitats. The significance was calculated using Wilcoxon’s rank-sum test.

Principal coordinate analysis (PCoA) based on the vOTU composition revealed a clear separation between the 3 habitats (Figure 1C). On the PCoA plot, gut virome was significantly stratified to the oral virome at the primary principal coordinate (PC1, explained 25.0% of total variance), while the dental plaque and saliva viromes had deviated at the secondary principal coordinate (PC2, explained 8.8% of total variance). Consistently, intuitive comparison of Bray-Curtis dissimilarity showed that the gut viromes were quite far away from the dental plaque and saliva viromes (Figure 1D).At the family level, all three habitats were dominated by viruses of five families (ignoring the family-level unclassified vOTUs): Siphoviridae, Myoviridae, Herelleviridae, Autographiviridae, and Phycodnaviridae (Figure 1E; Figure S3; and see Table S2 for the full list of viral families). The gut virome had a lower proportion of Siphoviridae and higher proportions of the other 4 families compared with dental plaque and saliva samples.

### Oral and gut viromes are disturbed in untreated and treated RA patients

Comparative analysis was conducted to investigate the differences in oral and gut viral communities among healthy controls and untreated and treated RA patients. Multiple comparisons were performed based on 1,944 high-abundance vOTUs with >0.01% mean relative abundance in all samples.

For dental plaque virome, the untreated and treated RA patients had a significantly lower Shannon index than the healthy controls (Figure 2A). For the viral richness, untreated patients were close to that of the healthy controls, while treated patients was still significantly lower than other subjects (Figure 2B). Distance-based redundancy analysis (dbRDA) revealed a visible deviation between three cohorts (Figure 2C). 57of 679 high abundant (>0.01% in all dental plaque samples)vOTUs had significantly differed in relative abundance among three cohorts (Table S3).These vOTUs included a *Lactococcus* phage (vOTU70) that enriched in the dental plaque virome of treated patients (Figure S4A); this phenomenon is potential linked to the usage of dairy products in some RA patients[31, 32].At the family level, four families, including Marseilleviridae, Ascoviridae, Microviridae, and Poxviridae, were differed in relative abundance among three cohorts (Figure 2D; Table S4).

**Figure 2.**
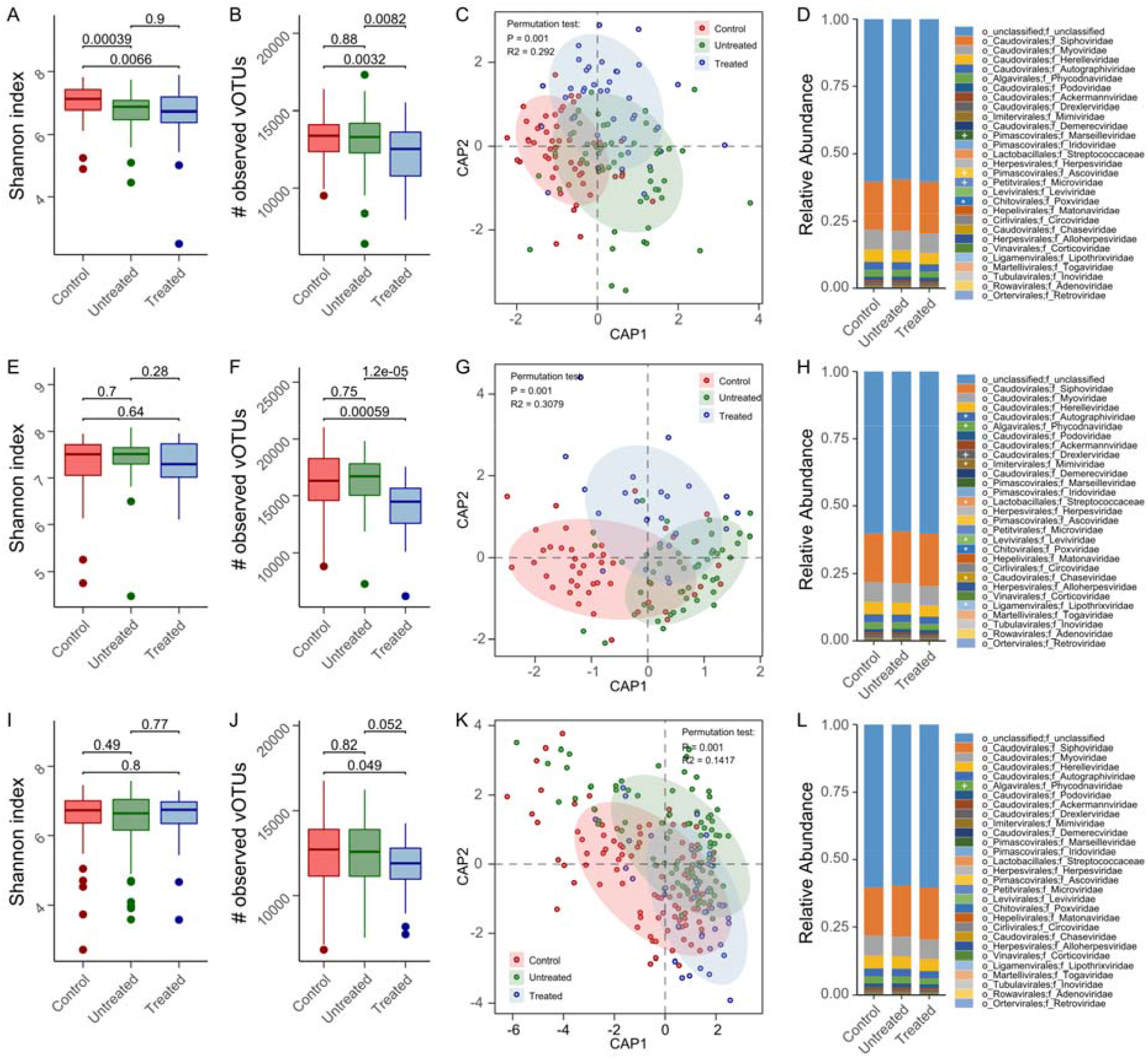
Community variation of the oral and gut viromes for RA patients. The alpha diversity indexes among healthy controls, untreated and treated patients with RA (**A-B** dental plaque, **E-F** saliva, **I-J** fecal). The significance was calculated using Wilcoxon’s rank-sum test. Distance-based redundancy analysis (dbRDA) based on Bray-curtis distance of the vOTU composition in healthy controls, untreated and treated patients with RA (**C** dental plaque, **G** saliva, **K** fecal). Family-level viral compositions in healthy controls, untreated and treated patients with RA (**D** dental plaque, **H** saliva, **L** fecal). The significance was calculated using the Kruskal-Wallis test. + adjusted *P* < 0.05, * adjusted *P* < 0.01.

For saliva virome, the Shannon index of the three cohorts was equal, whereas the viral richness of treated patients was significantly lower than that of untreated patients and healthy controls (Figure 2E-F).dbRDA revealed visible deviation between three cohorts (Figure 2G). 463 of 633 high abundant vOTUs had significantly differed in relative abundance among three cohorts (Table S5). Unlike the dental plaque virome, the *Lactococcus phage vOTU70* was enriched in the oral virome of both healthy subjects and treated RA patients (Figure S4B). At the family level, 9 families had differed in relative abundance among three cohorts (Figure 2H; Table S4),of which a dominant viral family, Autographiviridae, was significantly enriched in the untreated patients.

For gut virome, both the Shannon index and viral richness were similar in three cohorts (Figure 2I-J). dbRDA revealed visible deviation between three cohorts (Figure 2K), despite that the effect size of disease states was relatively small in gut virome (R^2^=14.2%) compared to dental plaque and saliva viromes (R^2^=29.2% and 30.8%, respectively). 73 of 975 high abundant vOTUs had significantly differed in relative abundance among three cohorts (Table S6). Besides, only the family Phycodnaviridae had differed among three cohorts (Figure 2L; Table S4), and it was significantly decreased in the gut virome of treated RA patients.

Taken together, our finding revealed that the viral diversity, vOTU numbers and family composition were remarkably altered in the oral virome of untreated and treated RA patients, while these alterations occurred at a relatively low incidence in the gut virome of RA patients.

### Interactions of viruses and bacteria are disrupted in treated RA patients

The bacterial microbiome (as well as the archaeal microbiome, hereinafter jointly referred to as “bacteriome”) of all samples was quantified by totaling 225 high abundant species (>0.05% in all samples) based on reads mapping of unique clade-specific bacterial/archaeal marker genes[33]. 44 (*P*<0.05, corresponding FDR 6.5%), 54 (*P*<0.05, corresponding FDR 3.4%), and 23 (*P*<0.05, corresponding FDR 20.4%) species had differed in relative abundance among three habitats (Table S7).

Speaking of the diversity, we found that the Shannon index was strongly correlated between virome and bacteriome in all three ecosystems and yet in all cohorts (Figure 3A; Figure S5), suggesting an extensive connection between viruses and bacteria. To study such interactions, we built the co-abundance networks between 75 high-abundance vOTUs and 225 bacterial species with an average abundance at>0.05% in all samples (Figure 3B).In dental plaque group, virus-bacterium associations amounted highest in the untreated RA patients, followed by the healthy controls, and RA patients came bottom (Figure 3C).Furthermore, healthy control and untreated RA patients were more closely associated than their respective with treated RA patients. Similar result was also observed in the gut virome (Figure 3E). In saliva, virus-bacterium associations appeared most in the healthy control cohort, while the treated RA patients still held a remarkable reduction (Figure 3D). Overall, these findings revealed that the viruses-bacteria interaction network was disrupted in treated RA patients.

**Figure 3.**
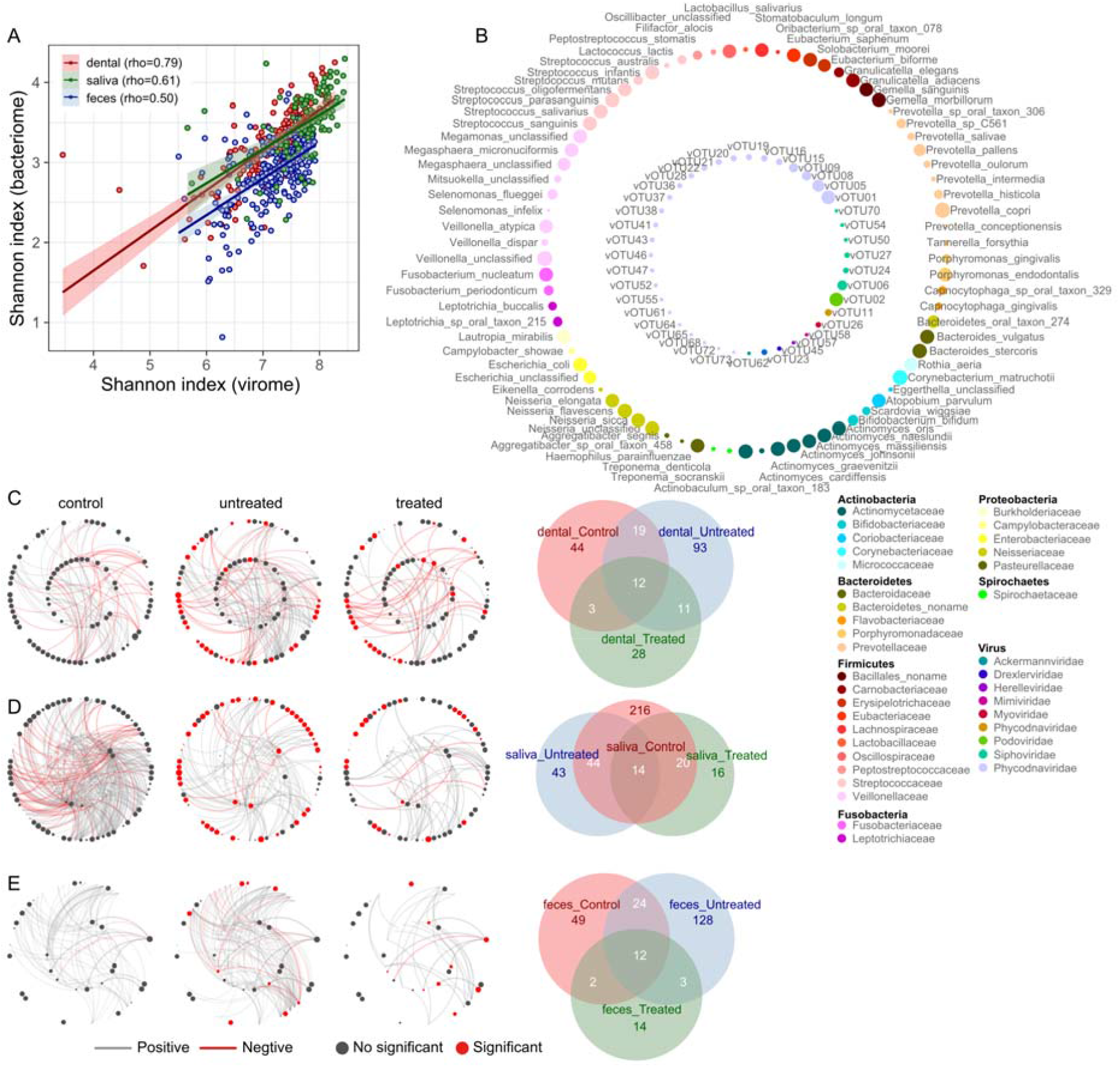
Correlation between bacteriome and virome. **A** The correlation analysis of Shannon index between bacteriome and virome in three habitats. **B** High abundant microbes with >0.05% average relative abundance in all samples. The size of node represented the mean relative abundance of vOTUs and bacteria species in all samples. **C-E** The correlation networks were based on the relative abundance of RA-associated vOTUs and bacteria species for healthy controls, untreated and treated patients (**C** dental plaque, **D** saliva, **E** fecal). Red node indicated microbes with significant difference (control vs patients, Wilcoxon’s rank-sum test, adjusted *P* < 0.05). Red line indicated a negative correlation, and grey line indicated a positive correlation (Spearman correlation test, adjusted *P* < 0.05). Venn diagrams showing the number of shared significant associations between RA-associated vOTUs and bacteria species in samples from healthy controls, untreated and treated patients.

### Oral viruses and bacteria are frequently linked to RA-associated clinical indexes

We then want to investigate the effects of host characteristics and clinical parameters on the oral and gut viromes and bacteriomes. In accordance with the aforementioned analysis, the stratification of three cohorts (untreated and treated RA status and healthy controls) significantly affected both oral and gut viromes and bacteriomes at the vOTU/species level, and this influence was greatest in saliva virome and bacteriome but slightest in gut virome and bacteriome(Figure 4A). Of the host demographic variables, age and gender had a considerable impact on the dental plaque virome and bacteriome but did not affect those of the gut, whereas the BMI and weight had a significant on the gut virome and bacteriome (Figure S6A).Of the clinical parameters, we found that several RA-associated indexes significantly impacted the oral virome. For example, CRP, GH, and RA_duration affected the dental plaque virome, while CDAI, DAS28, Disease_activity, and GH affected the oral virome (Figure 4B).Besides, oral virome were also influenced by several other clinical indexes, such as dental plaque virome being affected by HGB and saliva virome being affected by PLT and WBC (Figure S6B-C).

**Figure 4.**
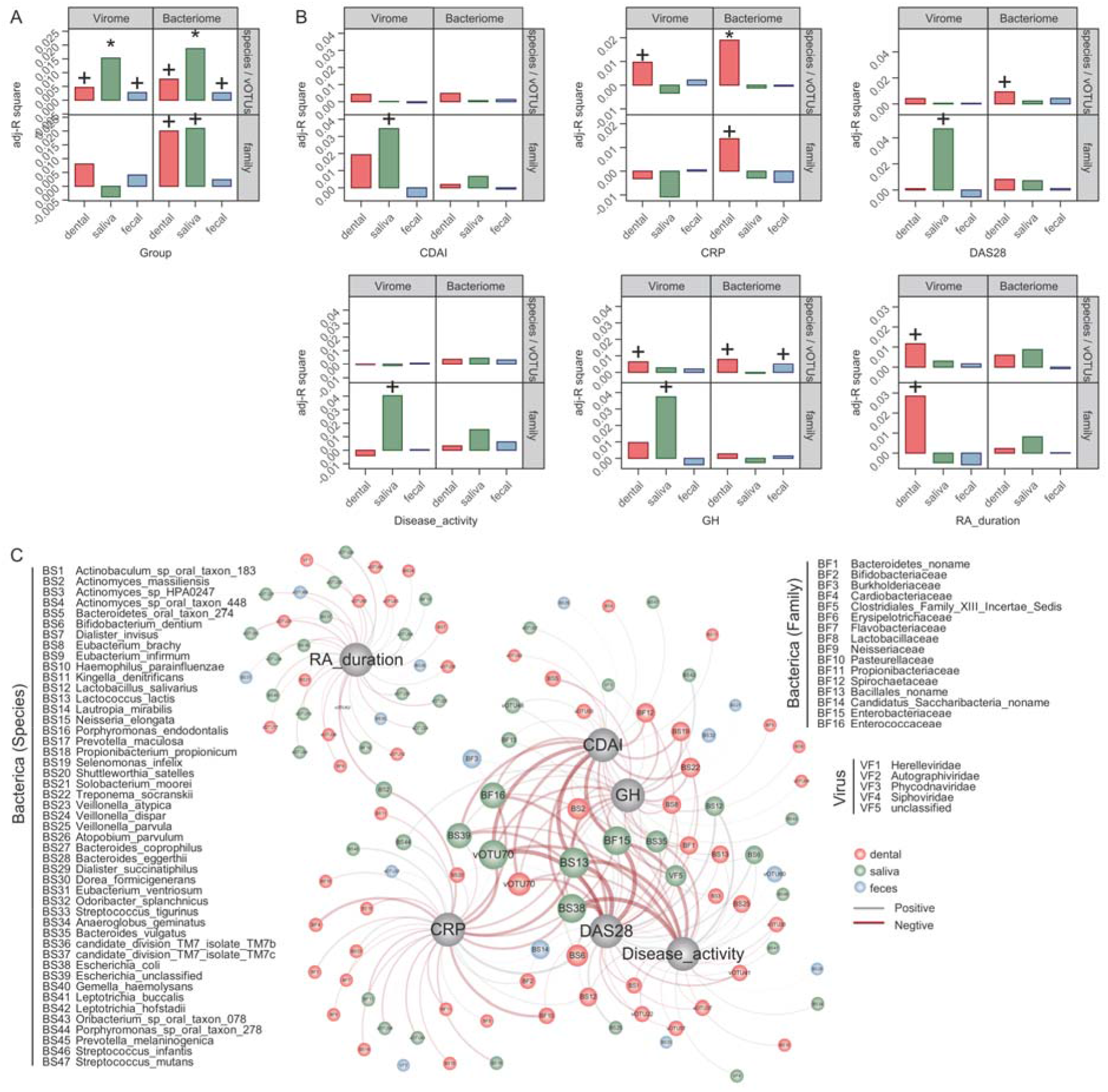
Correlation among the virome, bacterial microbiome and clinical indexes. **A** The effect size of RA status on the oral and gut virome and bacteriome (*Adonis*, + adjusted *P*<0.05, * adjusted *P*<0.01). **B** The effect size of several RA-associated indexes on the oral or gut virome. **C** The correlation network of 6 RA-associated clinical indexes and microbes (Spearman correlation test, adjusted *P* < 0.05). Edge weight indicated the strength of the correlation. Node size (for microbes) indicated the relative abundance of taxa in different habitats.

We generated a large correlation network of 6 RA-associated clinical indexes and microbes (including both viruses and bacteria with >0.05% average relative abundance in all samples) (Figure 4C).All indexes were correlated with a variety of bacteria and viruses from the dental plaque and saliva, but only a relatively small number of gut microbes were involved. Several dental plaque microbes, including *Actinomycesmassiliensis, Treponemasocranskii, Eubacteriumbrachy*, and *Lactococcus phage vOTU70*, as well as several saliva microbes, including *Lactococcus lactis, Bacteroides vulgatus, Escherichia coli*, and *Lactococcus phage vOTU70*, were the keystone taxa in the network, which were mainly associated with CDAI, GH, CRP, DAS28 and Disease_activity. To sum up, these findings revealed broad connection between oral viruses, bacteria and the RA-associated clinical indexes.

### Prediction of RA status employing viral and bacterial communities

Finally, we classified individuals of three statuses based on both bacteriome and virome using the random forest model. Each model featured with high abundant bacterial species and vOTUs (>0.05% relative abundance and significant difference). Each performance of model was assessed by the leave-one-out cross validation method. For dental plaque, the bacteriome achieved the AUCs of 80.54%, 84.57%, and 77.46% for discrimination of untreated patients vs. controls, treated patients vs. controls, and untreated vs. treated patients (Figure 5A), respectively. In dental plaque virome, the discriminability between untreated patients and controls was much lower (AUC=71.15%) than that between untreated vs. treated patients (AUC=94.3%) (Figure 5B). Combining bacteriome and virome achieved the AUCs of 81.57%, 89.4%, and 94.93% for discrimination of untreated patients vs. controls, treated patients vs. controls, and untreated vs. treated patients (Figure 5C), respectively. For saliva bacteriome and virome, the AUCs were nearly equivalentto that of dental plaque, with higher discriminability of untreated patients vs. controls and lower discriminability of treated patients vs. controls (Figure 5D-F). Inversely, for gut bacteriome and virome, the AUCs were significantly lower than that of dental plaque and saliva (Figure 5G-I). These results suggested that dental plaque and saliva performed high predictability for RA patients and healthy subjects, as well as for untreated and treated RA patients.

**Figure 5.**
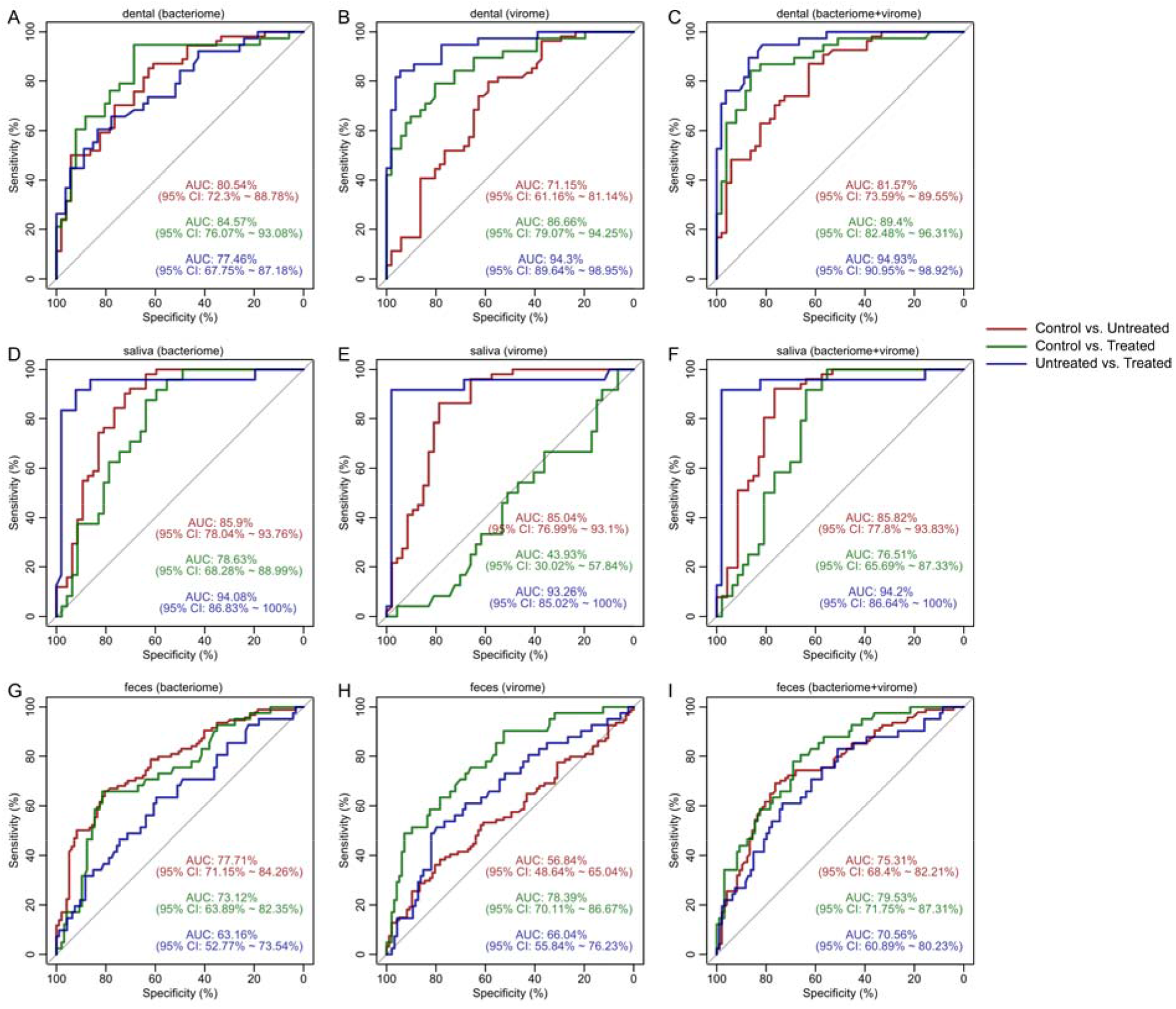
Classification of samples from healthy controls, untreated and treated patients with RA. RA-associated makers for fecal (**A-C**), dental (**D-F**), and salivary (**G-I**) samples from healthy controls, untreated and treated patients with RA were used to improved accuracy in the classification models.

## Discussion

We analyzed the oral and gut viral communities using the public dataset based on previous RA gut microbiome research[2]. Following the identification of 48,718 vOTUs, we discovered the alterations of oral and gut viral diversity and composition as well as virus-bacteria associations between untreated and treated RA patients. As some specific oral or other viruses have been reported to be associated with the etiology of autoimmune diseases in previous research[34, 35], this study further substantialized these viruses at metagenome-based virome level.

Oral habitats (dental plaque and saliva) had a lower proportion of virome sequences, but higher within-sample diversity compared with that of fecal samples, which was consistent with the previous findings that the human oral cavity had fewer bacteria but high diversity [36-38]. However, what the contrast was that the oral and gut bacterial contents are rarely shared (<5% sharing genes in Tierney *et al*’s study [38] and also <20% sharing genes in preliminary RA study [2]), and the oral and gut viromes are highly overlapped with >80% vOTUs in both habitats. We also found that the viral compositions of dental plaque, saliva, and gut samples were similar to some extent at the family level, despite of the highly-diverse bacterial composition [2]. These findings suggested that the oral and gut viruses might transfer or communicate more frequently than of bacteria. The potential mechanism of viral exchange between oral and gut viral communities needs further research to be illustrated.

Oral virome was more sensitive than other body sites to the environmental changes such as antibiotics taking [39]. In both untreated and treated RA patients, we found that the RA-associated changes of oral virome were remark ably greater than that of gut virome, so as in the bacterial metagenomic studies, suggesting that the oral microbiota could be the autoimmune regulator for its high correlation with host systemic autoimmune features[6, 40, 41]. Notably, in the oral virome, 57dental plaque and 463 saliva vOTUs had significantly differed in abundance among untreated/treated RA patients and healthy controls. One of these vOTUs, *Lactococcus phage vOTU70*, showed over growth in the dental plaque of treated RA patients and the saliva of both controls and treated patients. This finding indicated the potential connection of *Lactococcus* phages and RA treatment. With relation to these variations in viral composition between cohorts, the oral virome were more sensitive to the RA-associated clinical parameters. For example, the saliva virome could be significantly influenced by two parameters regarding RA disease activity (DAS28 score and Disease_activity), suggesting it maybe potential indicator. This finding complied with recent opinions, that the RA patients’ oral health condition was connected with their disease activity[42]. Also, the dental plaque virome was influenced by CRP and RA_duration, suggesting oral viruses played a role both for the short-term inflammation level and the long-term disease progression in RA patients.

Effects of disease status on gut virome was considerable, as 73 vOTUs were identified with different abundance among untreated, treated patients and controls. Moreover, the co-altered oral and gut viruses found in this study suggested a potential oral-gut virome axis in the pathogenesis and treatment progresses of RA, which was partly consistent with the recent studies proposing the oral-gut microbiome axis in the etiology of RA [43, 44]. Some studies had shown that the Epstein-Barr (Eb) virus could cause RA by infecting synovial tissues[45, 46], however, this virus was not identified in our oral and gut viromes dataset.

Close interactions of gut virome and bacterial microbiome had been reported in patients of several diseases such as inflammatory bowel disease (IBD)[17, 47] and cirrhosis[48]. Here, we enriched their relationships in the oral ecosystem by constructing the virus-bacteria co-abundance networks in both dental plaque and saliva samples. Interestingly, we found that the virus-bacteria network in both untreated and treated RA patients was remarkedly changed compared with that in healthy controls, and it was much more considerable in treated RA patients, with the manifestation as virus-bacteria association decrease in all three body sites. The alteration of virus-bacteria interactions might suggest a perturbance of the whole ecosystem[49], establishing deeper connection with the pathogenicity of RA.

Finally, we explored the feasibility of combining bacteriome and virome for the discrimination of RA and healthy controls. Results showed that AUC could achieve 89.4% in discriminating controls and treated patients (in dental plaque samples), 85.82% in discriminating controls and untreated patients (in saliva samples), and 94.2% in discriminating treated and untreated patients (in saliva samples), respectively. These AUC values were equal to or larger than that based on bacteriome or virome alone, indicating that the virome is effective for distinguishing RA status and for early diagnosis, despite systematic investigations of key viral markers might be helpful in the future. A similar predictable effect of virome was also observed in the gut of IBD patients [17].

## Methods

### Virus identification and analyses

#### Metagenomic data and assembly

The metagenomic data of 497 samples were obtained from NCBI with project accession no. PRJEB6997.The quality control of metagenomic reads was performed using fastp[50], and the human reads were removed based on Bowtie2 alignment [51]. Each sample was individually assembled using metaSPAdes[52]. Proteins of the assembled contigs were predicted using Prodigal[53].

#### Viral identification and vOTUs

After assembly, the assembled contigs(≥5,000 bp) were identified as viruses when they satisfied one of the following criteria: 1) at least 50% proteins of a contig (or at least 3 proteins if the contig had less than 6 proteins) were assigned into the viral protein database integrating from NCBI reference viral genomes and the virus orthologous groups database (http://vogdb.org), with a maximum pairwise alignment e-value 1e-10 based on DIAMOND[54]; 2) classified as categories 1, 2, 4, and 5 viruses in the VirSorter[55], a homology-based viral identifier from assembled metagenomic data; and 3) score >0.9 and p-value <0.05 in the VirFinder[56], a k-mer based tool for identifying viral sequences. Viral contigs were pairwise blasted and the highly consistent viruses with 95% nucleotide identity and 80% coverage of the sequence were further clustered into vOTUs using in-house scripts. The longest viral contig was defined as a representative sequence for each vOTU. Proteins of the vOTUs were aligned with the available viral proteins using BLASTP (minimum score 50), and the family level taxonomy of a vOTU was generated if more than a third of its proteins were assigned into the same viral family. Lastly, the genome quality of vOTUs was assessed by CheckV[30].

#### Taxonomic abundances

To calculate the relative abundances of vOTUs, clean reads in each sample were first mapped against vOTU contigs using Bowtie2 alignment. Reads mapped to each vOTU were aggregated using SAMtools[57]. The relative abundances of each vOTU were the total number of assigned reads divided by the total number of reads mapped to all vOTU in each sample. The relative abundance profiles of viral families were generated by adding up the abundance of vOTUs annotated to the same family.

### Bioinformatic and statistical analyses

#### Alpha and beta diversity

Rarefaction analysis was performed to estimate sequencing depth using the custom script. Alpha diversities in each sample were estimated based on the rarefaction data: 1) the number of observed OTUs were the number of vOTUs with non-zero abundance; 2) Shannon diversity index was calculated using the *vegan*’s diversity function. Beta diversity was evaluated using Bray-Curtis dissimilarity based on the relative abundance of vOTUs.

#### Multivariate statistics

All statistics and figure generation were carried out in R (v. 4.0.2).Principal coordinate analysis (PCoA) was performed based on the relative abundance profile of vOTUs using the function *dudi*.*pcoa* from the R package *ade4*. Distance-based redundancy analysis (dbRDA) based on Bray-Curtis dissimilarity was carried out using the *capscale* function of the R package *vegan*, and the significance value was estimated based on 1,000 permutations by the function *envfit*. In addition, to evaluate the explained variation of the different covariate on the microbial composition, Permutational multivariate analysis of variance (PERMANOVA) was performed using the function *adonis*. The effect size of a variable was determined as R-square value adjusted by the function *RsquareAdj*, and p-value < 0.05 was considered significant.

#### Prediction model

Random forest models were carried out based on vOTUs or bacterial species with a significant difference among disease groups, using the function *randomForest*. ROC curves with leave-one-out cross-validation were used to evaluate the model using the *roc* function of the R package *pROC*.

#### Statistical test

To determine associations between taxa and covariates or between vOTUs and bacterial species, Spearman correlation coefficient was measured using the function *cor*.*test* in R.Significance tests between the two groups for taxonomic abundances were performed using the function *wilcox*.*test*, while significance tests among multiple groups were performed using the function *kw*.*test. P*-values fromthe comparative analysis were adjusted using the function *fdrtool. P*-values from correlation analysis were adjusted using the function *p*.*adjust* with the ‘method=BH’ parameter.A *P*-values<0.05 was considered different and an adjusted*P*<0.05 was considered significant.

## Conclusions

Our results demonstrated community variation among dental plaque, saliva and feces viromes. In oral and gut samples from untreated and treated RA patients, the perturbance of viral composition and correlation network between microbes and RA-associated clinical indexes might be concerned with the pathogenicity of RA.

## Acknowledgements/funding

This work was supported by grants from the Shenzhen Puensum Genetech Fund (No. 20200106), the National Natural Science Foundation of China (No. 81902037), and Dalian Science and Technology Innovation Fund(2020JJ27SN069).

---

**Figure S1.**
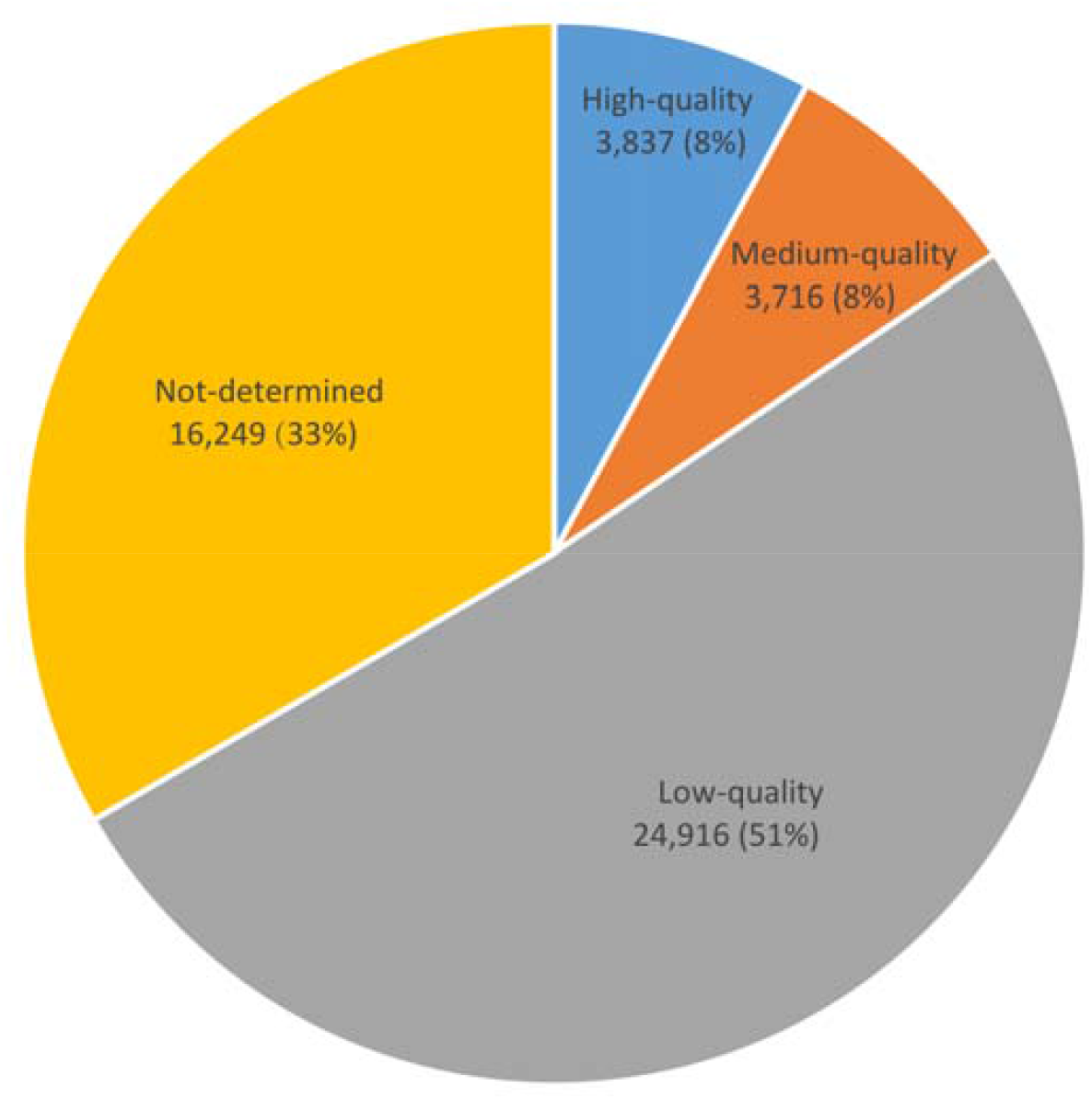
ChevkV assessment of 48,718 vOTUs.

**Figure S2.**
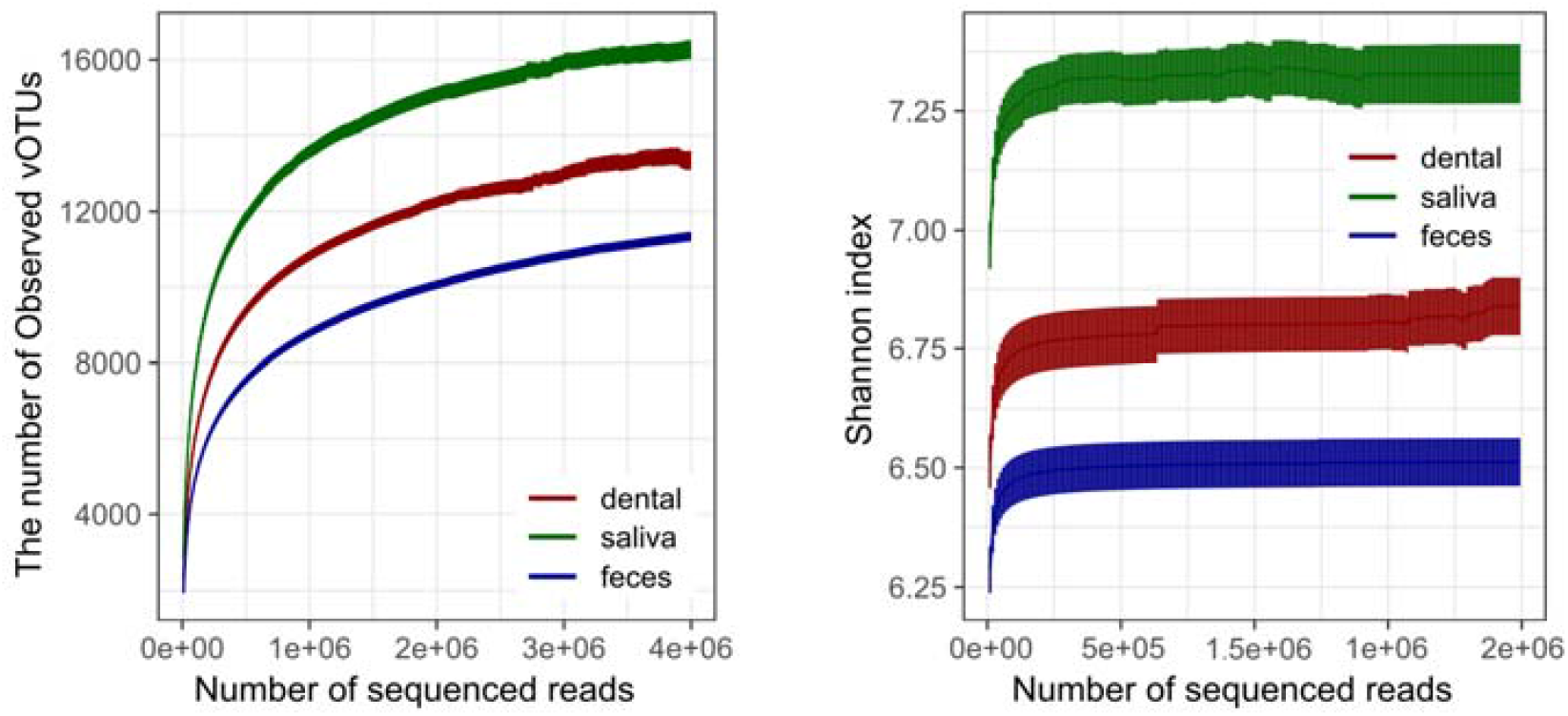
Rarefaction curves for the number of observed vOTUs and Shannon index.

**Figure S3.**
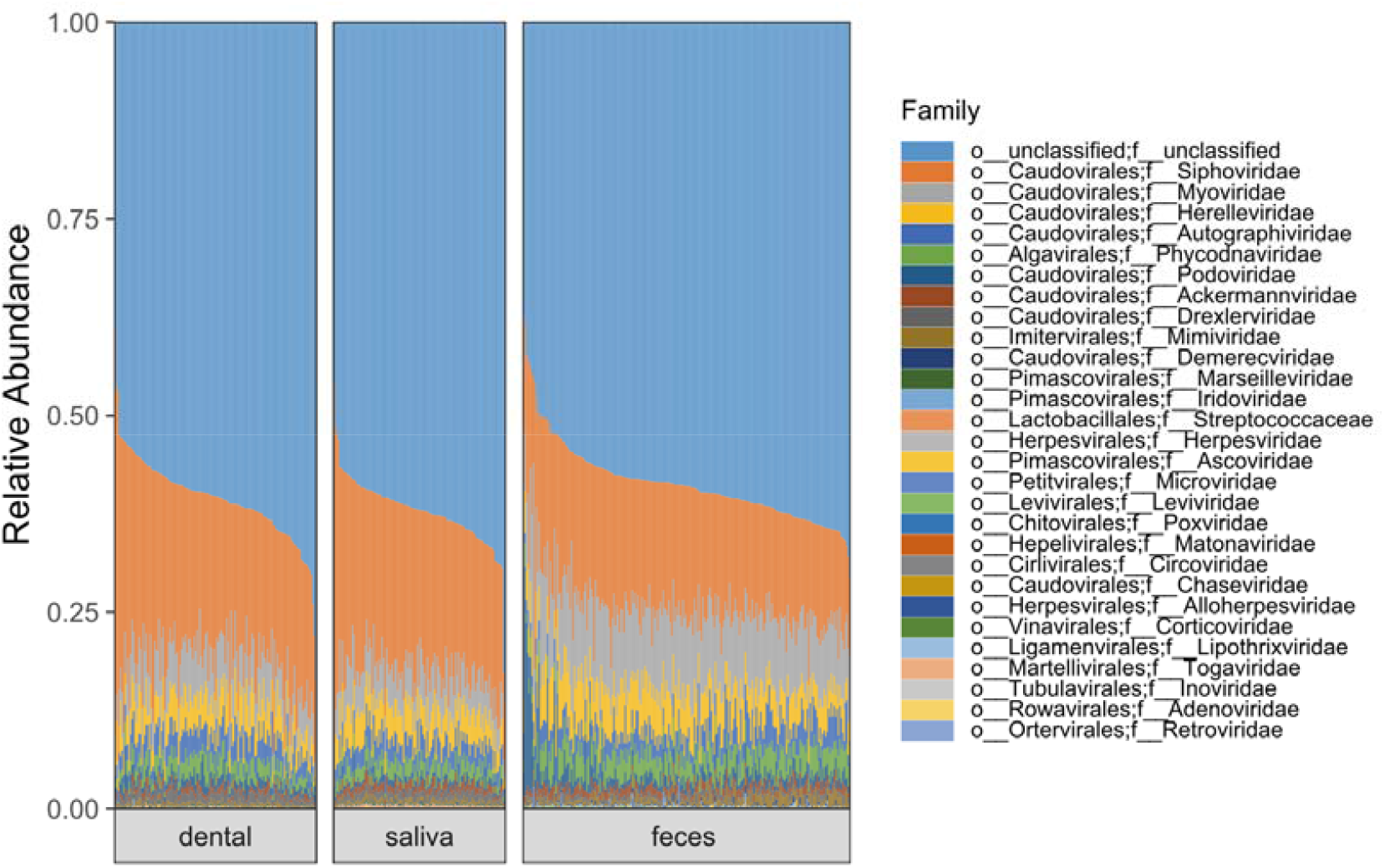
Viral composition at the family level in all dental plaque, salivary, and fecal samples.

**Figure S4.**
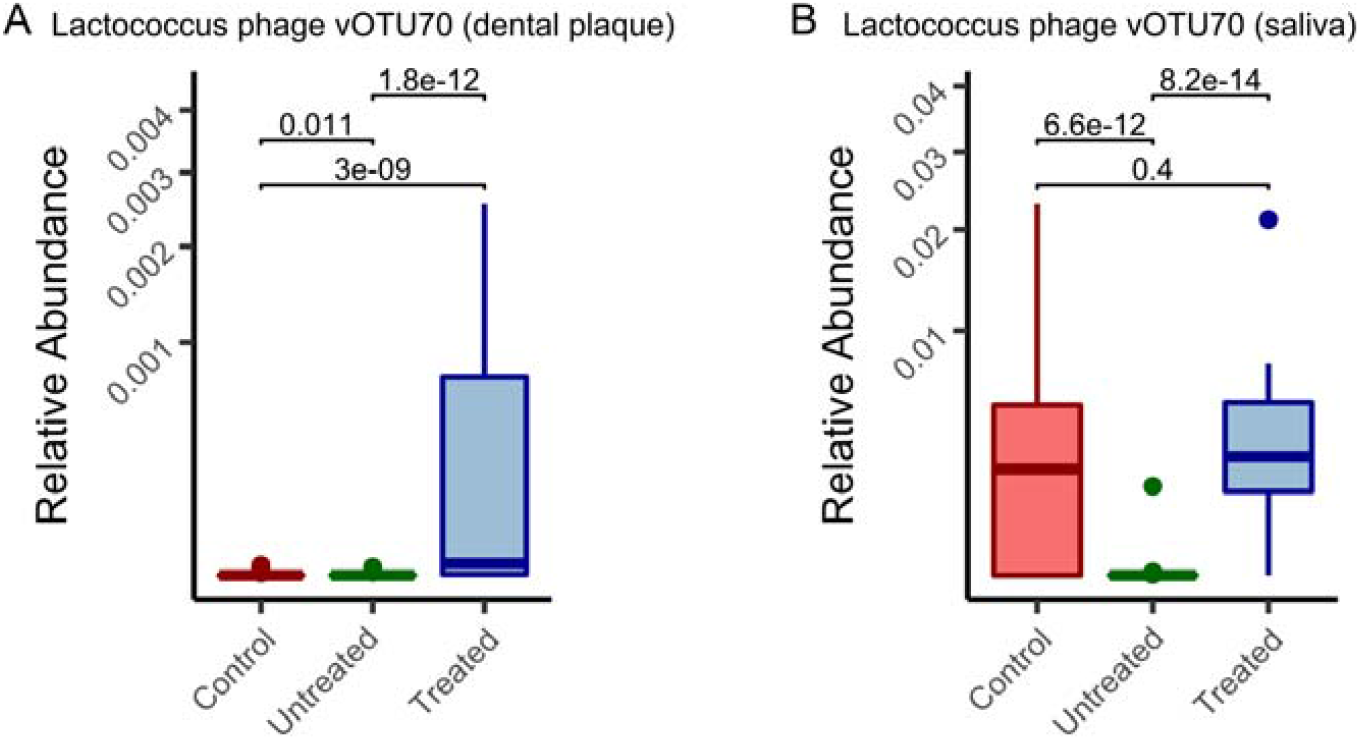
Comparison of relative abundance of *Lactococcus phage vOTU70* among healthy controls, untreated and treated patients with RA. A Dental plaque samples. **B** Saliva samples.

**Figure S5.**
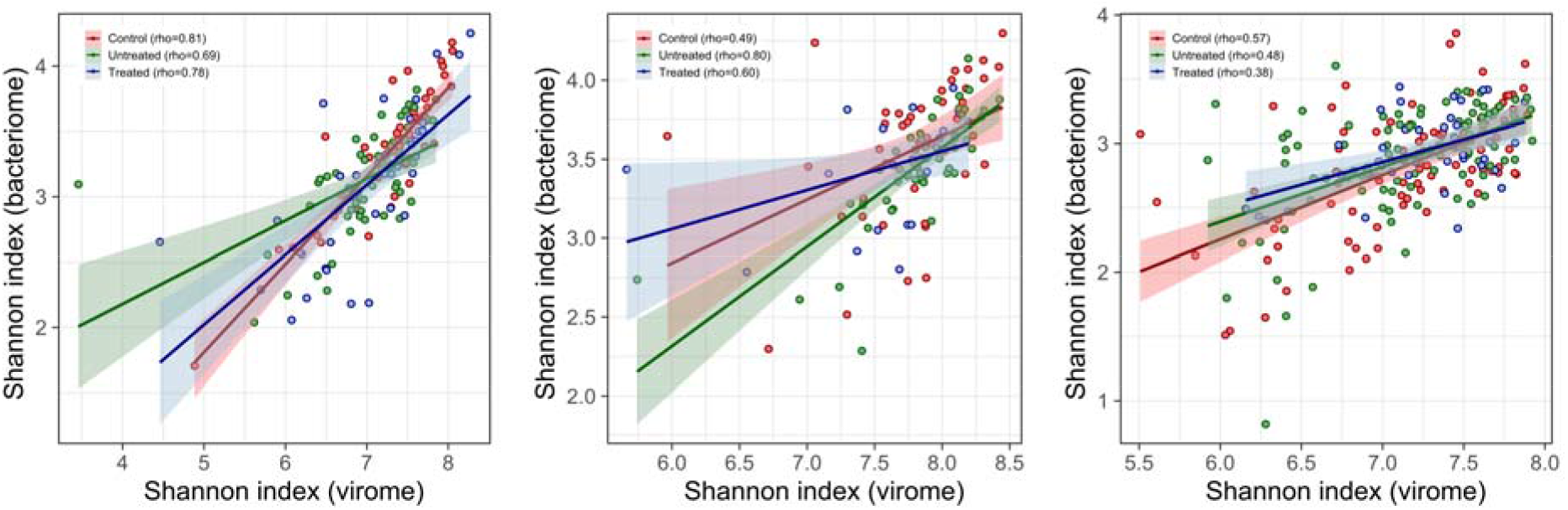
The correlation analysis of Shannon index between bacteriome and virome in oral and gut samples from healthy controls, untreated and treated patients with RA. Left plot, dental plaque. Center plot, saliva. Right plot, feces.

**Figure S6.**
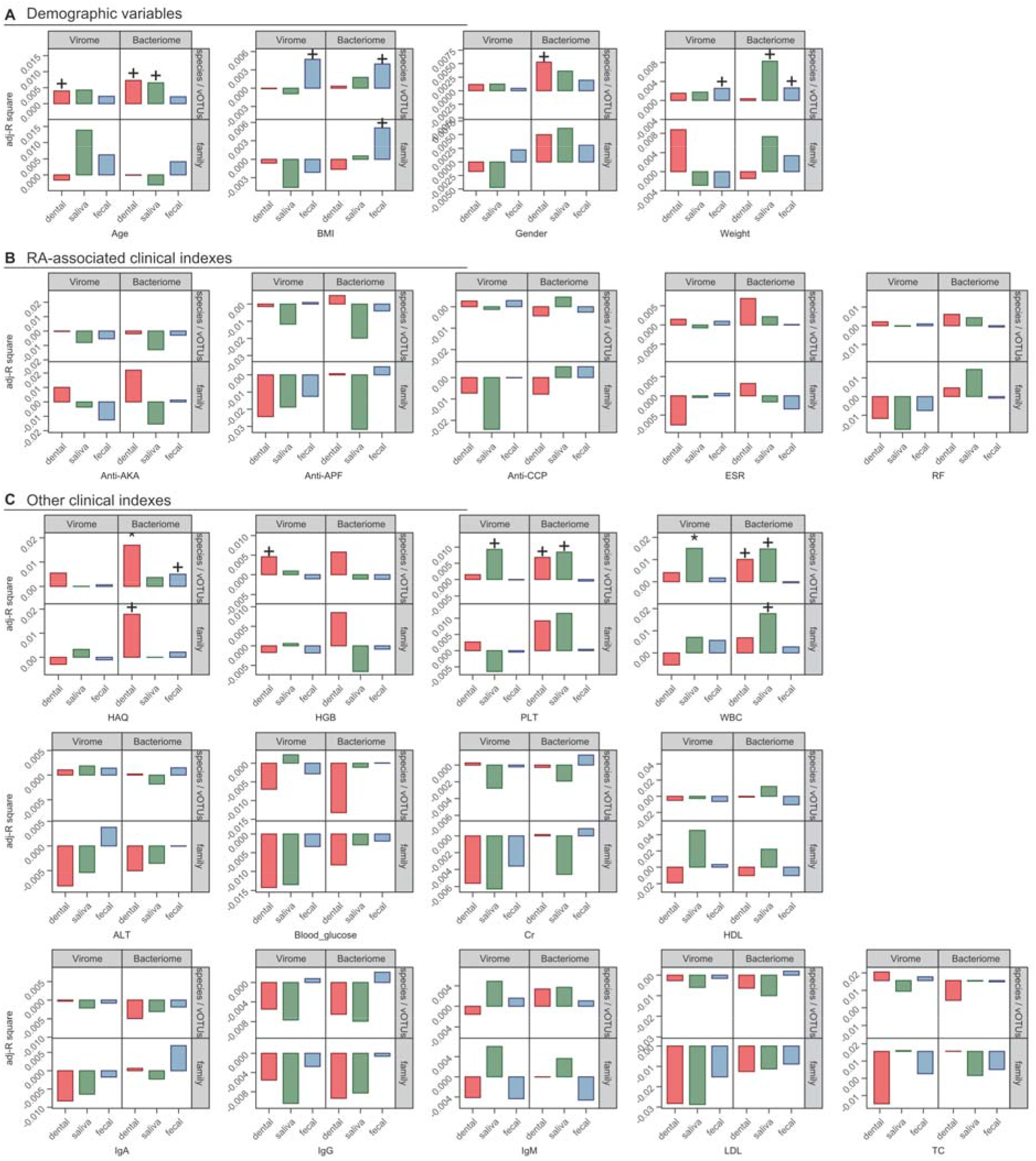
The effect size of clinical indexes on the oral and gut virome and bacteriome. *Adonis*, + adjusted *P*<0.05, * adjusted *P*<0.01.

